# Reaction of ependymal cells to spinal cord injury: a potential role for oncostatin pathway and microglial cells

**DOI:** 10.1101/2021.02.12.428106

**Authors:** R. Chevreau, H Ghazale, C Ripoll, C Chalfouh, Q Delarue, A.L. Hemonnot-Girard, H Hirbec, S Wahane, F Perrin, H Noristani, N Guerout, JP Hugnot

## Abstract

Ependymal cells with stem cell properties reside in the adult spinal cord around the central canal. They rapidly activate and proliferate after spinal cord injury, constituting a source of new cells. They produce neurons and glial cells in lower vertebrates but they mainly generate glial cells in mammals. The mechanisms underlying their activation and their glial-biased differentiation in mammals remain ill-defined. This represents an obstacle to control these cells. We addressed this issue using RNA profiling of ependymal cells before and after injury. We found that these cells activate STAT3 and ERK/MAPK signaling during injury and downregulate cilia-associated genes and FOXJ1, a central transcription factor in ciliogenesis. Conversely, they upregulate 510 genes, six of them more than 20 fold, namely *Crym, Ecm1, Ifi202b, Nupr1, Rbp1, Thbs2 and Osmr*. OSMR is the receptor for the inflammatory cytokine oncostatin (OSM) and we studied its regulation and role using neurospheres derived from ependymal cells. We found that OSM induces strong OSMR and p-STAT3 expression together with proliferation reduction and astrocytic differentiation. Conversely, production of oligodendrocyte-lineage OLIG1^+^ cells was reduced. OSM is specifically expressed by microglial cells and was strongly upregulated after injury. We observed microglial cells apposed to ependymal cells in vivo and co-cultures experiments showed that these cells upregulate OSMR in neurosphere cells. Collectively, these results support the notion that microglial cells and OSMR/OSM pathway regulate ependymal cells in injury. In addition, the generated high throughput data provides a unique molecular resource to study how ependymal cell react to spinal cord lesion.

## Introduction

The spinal cord lies in the caudal part of the central nervous system and conveys motor information to the muscles and sensory signals back to the brain. It is affected by several pathologies such as multiple sclerosis, motoneuron degeneration and traumatic injuries. No curative treatments for these diseases exist. Animals like salamanders and zebrafish regenerate spinal cord cells, including neurons, after lesions (Becker, Becker, et Hugnot 2018). This extraordinary property is due to the persistence in adults of fetal-like radial-glia stem cells around the spinal cord central canal. These cells express the FOXJ1 transcription factor and keep expressing spinal cord developmental genes such as *Pax6* and *Shh*. Similar FOXJ1^+^ cells exist in mammals, including young human, constituting the ependymal region ((Marichal et al. 2017; Hamilton et al. 2009; Sabelström, Stenudd, et Frisén 2014; Garcia-Ovejero et al. 2018). We and others have shown that this region is organized like the germinative cell layer (namely the neuroepithelium) which generates most spinal cord cells during development. Key spinal cord developmental transcription factors and genes such as *Arx, Msx1, Nestin, Pax6, Zeb1* remain expressed in this region (Ghazale et al. 2019).

Mouse ependymal cells are ciliated and hardly proliferate in the normal situation. However, it is known since 1962 (Adrian et Walker 1962) that after spinal cord injury (SCI) they resume proliferation under the influence of Ras signaling (Sabelström et al. 2013) and then migrate to the lesion site. This is principally observed when the lesion compromises the integrity of the ependymal region (Ren et al. 2017). However, unfortunately, they produce no neurons and few oligodendrocytes and mainly form astrocytes (Barnabé-Heider et Frisén 2008). These astrocytes contribute to the core of the glial scar and are beneficial for recovery (Sabelström et al. 2013). Importantly, when grafted in a neurogenic environment (hippocampus), cultured spinal cord ependymal cells generate neurons (Shihabuddin et al. 2000). Also in vitro, a fraction of ependymal cells form neurospheres which can be propagated for several passages and then induced to differentiate into neurons and glial cells (Barnabé-Heider et al. 2010). This shows that at least some ependymal cells are multipotent and depending on the context, can form new neurons or glial cells. By manipulating these cells in vivo by expression of oligodendrogenic transcription factors, it was recently shown that they can replace large numbers of lost oligodendrocytes in the injured mouse spinal cord (Llorens-Bobadilla et al. 2020).

Theoretically, such plastic cells represent an attractive endogenous source to alleviate various spinal cord lesions if one could control their proliferation and fate to produce new neurons and oligodendrocytes. However, the molecular events taking place in these cells after SCI remain still largely unknown. This lack of knowledge makes it difficult to control these cells. Notably, we need to know: which genes are modulated in ependymal cells after injury ? which signaling is activated ? what is the influence of parenchymal spinal cord cells, notably microglial cells, on ependymal cell fate ? why ependymal cells mainly generate astrocyte after lesion ? The present work was designed to address these key issues.

Here we determined the RNA profiling of these cells before and after lesion. Ependymal cells are difficult to purify in vitro and their enzymatic dissociation could lead to artifactual RNA variations. To overcome this obstacle, the ependymal region of unfixed spinal cord sections was directly microdissected with a laser. This led us to uncover the implication of several signaling and genes. We used immunofluorescence and cell cultures to study these results further in vitro. Notably we discovered a role for the OSM/OSMR pathway and microglial cells in regulating ependymal cell properties. This new molecular resource and new knowledge will help to understand properties of these cells after injury.

## Material and Methods

### Animals

Animals were handled at the animal care facility in compliance with the Committee of the National Institute of Health and Medical Research (INSERM) who approved this study in accordance with the European Council directive (2010/63/UE) for the protection and use of vertebrate animals. All animals were handled under pathogen-free conditions and fed chow diet ad libitum. Adult CD1 mice (3 months, Charles River, France) were used for microdissection after spinal cord injury, RNA profiling, neurosphere cultures and histology. C57BL/6 mice aged 3-4 months were used for spinal cord organotypic slice cultures and histology. CX3CR1^+/GFP^ C57BL/6 (Jung et al. 2000) (Jackson Laboratories) transgenic mice aged 3 months were used for spinal cord microglial cell histology. β-actin-GFP C57BL/6 mice (Okabe et al. 1997) (Jackson Laboratories) aged 3 months were used to derive GFP^+^ neurospheres to co-culture with microglia BV-2 cells. For neurosphere cultures, RNA and protein extractions, adult spinal cords were dissected from mice euthanized by intraperitoneal injection of sodium pentobarbital (100 mg/kg).

### Spinal cord injury

Adult CD1 mice aged 3 months were anesthetized using isoflurane gas (1.5%). T9–T10 laminectomy was performed to expose the spinal cord. Four needle penetrations (30G) were done (staggered holes, 2 on each side of the posterior spinal vein). Penetration depth was approximately 1 mm. Muscles and skin were sutured and animal placed on heated pads until they wake-up. For control experiments, we performed laminectomy but no needle insertion (sham injury).

### Histology and immunofluorescence (IF)

For spinal cord histology, mice were anesthetized by intraperitoneal injection of sodium pentobarbital (100 mg/kg) and perfused intracardially with 10 mL of PBS followed by 50 mL of 4% formaldehyde-PBS solution (pH 7.0). After dissection, spinal cords were post-fixed in the same solution for 1 h at 4°C and cryopreserved by successive immersion in 10%, 20%, and 30% sucrose solutions in PBS for at least 6 h. Thoracic parts of the spinal cord were cut, embedded in OCT medium, rapidly frozen in liquid N2-cooled isopentane, and cryosectioned (14 µm) (Leica apparatus). IF were performed on slide-mounted spinal cord sections with primary antibodies and dilutions listed in supplemental information table S1 after permeabilization for one hour with 0.1-0.3 % Triton 100x and 5-10% donkey serum. For cell cultures, IF were done on cells grown on poly-D-lysin/laminin coated glass-coverslips fixed with paraformaldehyde (4%, 20 min, room temperature). Secondary antibodies (Alexa488 or Alexa594-conjugated specie-specific anti mouse, rabbit or goat) were purchased from the Jackson Laboratories. Incubations without primary antibody or with antibody recognizing antigens not present in the sections (monoclonal anti DYKDDDDK Tag or polyclonal anti GFP) were used as negative controls. Nuclei (blue in all images) were stained with Hoechst 1 µg/ml for 10 min. For p-STAT3 staining, antigen retrieval was done for 25 minutes at 90°C in citrate buffer pH 6.0 in a histology microwave oven (Histos 5, Milestone). The quality of stainings was evaluated by two independent investigators (JPH,HG, RC or CR). For stainings on SCI sections (Fig. 2), we selected sections where the needle tract was present to make sure we were close to the injury point. Luxol fast blue and neutral red staining (Fig. 1B) was performed as described in (Lockard et Reers 1962). All presented images are representative images and the number of examined sections and animals are indicated in the legends.

**Figure 1:**
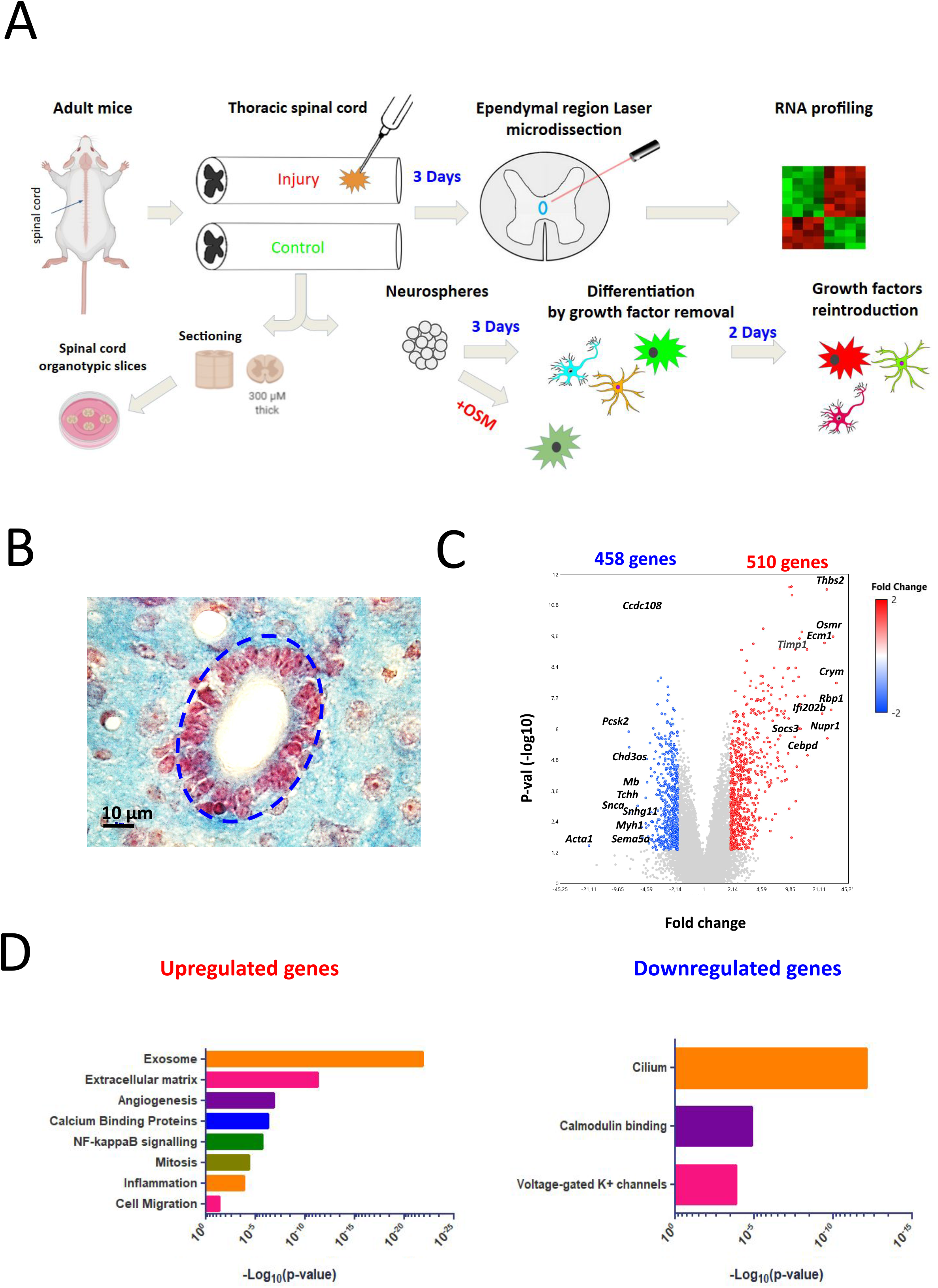
Bioinformatics analysis of ependymal gene expression in SCI. A: Schematic overview of main article steps and techniques. **B:** Mouse spinal cord ependymal region. Ependymal cell nuclei (thoraco-lumbar level) are stained in red with neutral red dye. Surrounding parenchyma is stained in blue with luxol fast blue. The microdissected area is delimited with blue dotted circles. **C:** Volcano plots of genes whose expression are dysregulated after injury in the ependymal region (fold change ≥ 2 or ≤ −2). Top10 up and downregulated genes are indicated. **D:** Selection of Gene Ontology analysis for up-and down-regulated genes with p-values. Comprehensive gene ontology analysis is provided on Table S2.

**Figure 2:**
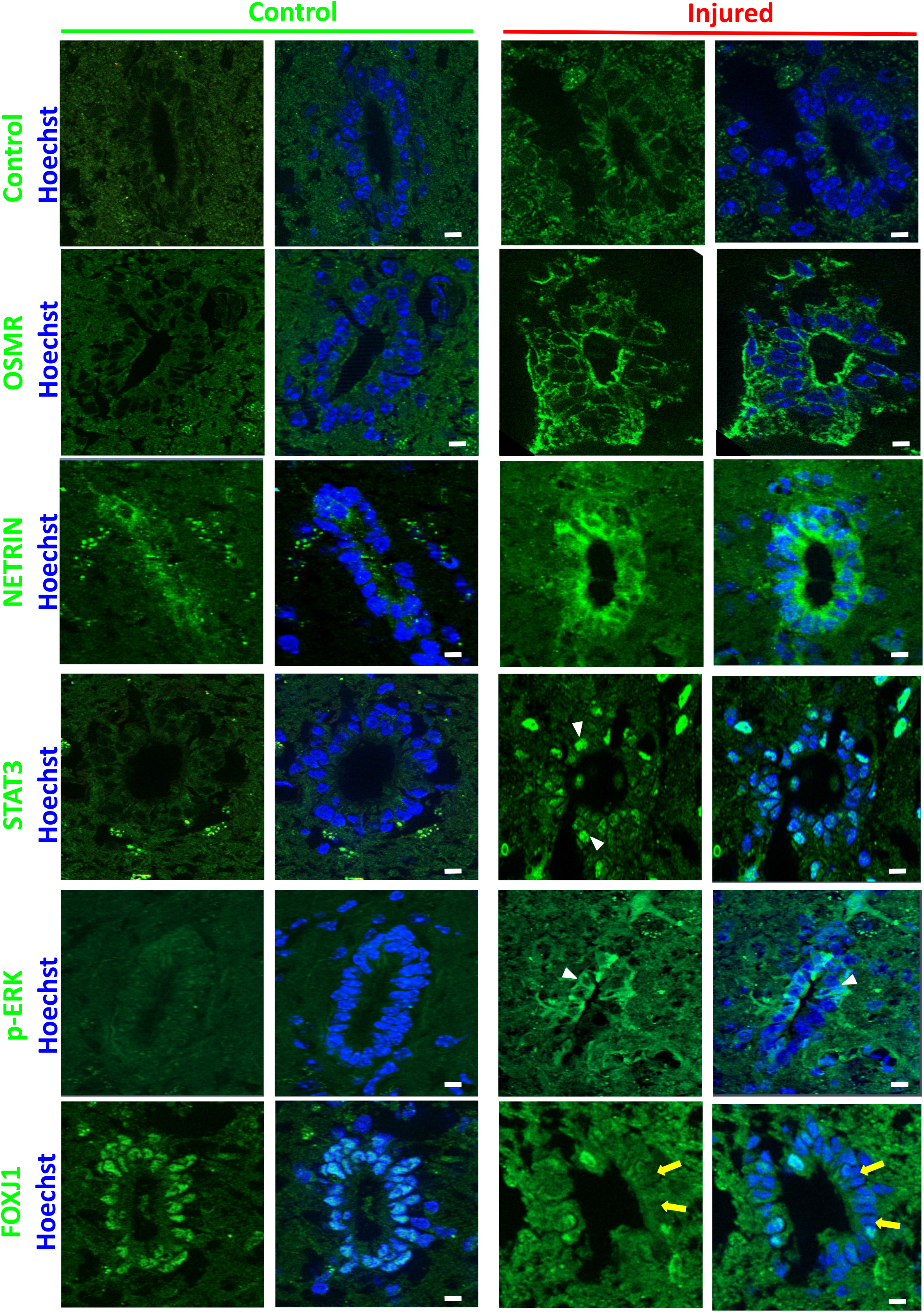
Immunofluorescence validation. Immunofluorescences performed on control and injured spinal cord for indicated proteins. Control immunofluorescence was performed using antibody against the synthetic DYKDDDDK TAG antigen. White arrowheads and yellow arrows show examples of positive and negative cells respectively. These images are representative of 2 independent experiments (n=6 animals in total, 7-15 sections examined per animals). Scale bars=10 µm.

### RNA extraction, RNA profilings and bioinformatics analysis

Laser microdissection of the ependymal region in sham and injured spinal cord (72h post injury, 6 animals each condition) were performed exactly as described in details in (Ghazale et al. 2019). After dissection, part of the thoracic spinal cord containing the needle insertion sites was selected and flashed frozen in N2 without chemical fixation. RNA from laser-microdissected areas were extracted with a ReliaPrep RNA cell Miniprep kit (Promega, Madison, USA) according to the manufacturer’s protocol and quantified/qualified with Nanodrop and Agilent 2100 Bioanalyzer apparatus (RNA Integrity Number (RIN) > 8.0). RNA labelling and profiling was performed using Affymetrix mouse microarrays as described in (Ghazale et al. 2019). Gene expression data were normalized with RMA algorithm and analyzed using the Affymetrix TAC 4.0 software (Transcriptome Analysis Console). The filter criteria were set to a linear fold change ≥ 2 or ≤ −2 (control vs injured spinal cord) with p value ≤ 0.05. Gene lists were analyzed with DAVID Bioinformatics Resources 6.83 for gene enrichment analysis (Huang, Sherman, et Lempicki 2009). Data were also processed to search for pathway activation (Fig. S1B) through the use of ingenuity pathway-analysis (IPA) software (Qiagen Inc.) (Krämer et al. 2014).

### Cell cultures

Neurosphere cultures were derived from adult spinal cord using the protocol and medium detailed in (Hugnot 2013). In the growing condition, medium contained EGF, FGF2 (10 ng/ml each, Peprotech) and Heparin (2 µg/ml, Sigma H3149). For differentiation, enzymatically-dissociated neurospheres were plated on poly-D-lysine/Laminin (1 µg/cm^2^) coated glass coverslips in a medium without growth factors and heparin but containing 2% fetal bovine serum (Invitrogen). After 4 days, the coverslips were either fixed for immunofluorescence or processed for RNA extraction (ReliaPrep RNA Miniprep kit, Promega) for microarray analysis. To test the influence of cytokines on cell growth/differentiation and OSMR expression (QPCR, Fig. 5C), all cytokines (Peprotech, USA) were used at 10 ng/ml. Influence of OSM on cell growth (Fig. 5G) was measured by seeding dissociated cells (5000 cells per well, 6 wells) in 1 ml of media in 24-well plates coated with poly-HEMA (Sigma P3932) to inhibit cell adherence. After 5 days, the neurospheres were directly dissociated by addition of trypsin in the wells (0.5% final) and the cell number was measured with an automated cell counter (Z2, Beckman Coulter).

To assess the influence of microglia on spinal cord stem cells, BV-2 immortalized microglial cells (Blasi et al. 1990) (passage 3 to 8) cultured in macrophage serum-free medium (ThermoFisher 12065-074) were used. Neurosphere cells derived from β-actin GFP spinal cord were cultured alone or in the presence of BV-2 cells (ratio BV-2 vs spinal cord cells was 1:6) in the neurosphere medium containing growth factors. After 3 days, cultures were dissociated with trypsin/EDTA 0.5% and GFP^+^ spinal cord cells were collected by FACS for RNA extraction and QPCR.

For spinal cord organotypic slices, the protocol described in (Fernandez-Zafra, Codeluppi, et Uhlén 2017) was used except that glutamine (2 mM final concentration) was used instead of glutamax and poly-D-lysine instead of poly-L-lysine for coating. Slices (300 µm) were fixed with paraformaldehyde (4%, 20 min, room temperature) directly after sectioning (t=0) or after 72 h in culture. Slices were then cryopreserved in PBS-sucrose 10, 20, 30% and embedded in OCT for cryosectioning and immunofluorescence.

### QPCR

cDNA synthesis was performed using 1-5 μg of total RNA with random hexamers and reverse transcriptase (Promega, GoScript). Quantitative RT-QPCR was performed in triplicate for each samples using the KAPA SYBR PCR kit (Sigma Aldrich KK4600) with a LightCycler 480 apparatus (Roche). 1 μL of cDNA was used for each reaction. Primers are listed in table S1. Relative expression values were calculated using the 2-ΔΔCT method and normalized using the β-actin gene. All QPCR reactions were performed with three independent cultures.

### Western blot

Total proteins from spinal cord samples and cultured cells were extracted and used for Western blots as described in (Azar et al. 2018). Proteins were detected using the Odyssey CLx Li-Cor technology. Briefly, primary antibodies were incubated in Li-Cor PBS buffer overnight at 4°C. After washing, membranes were incubated with secondary fluorescent dye (IRDye 800CW for OSMR protein and IRDye 680LT for β-actin normalization). For figure 5D-F, signals were obtained with peroxidase-conjugated secondary antibodies (Cell Signalling Technology) and revealed with Clarity Western ECL kit (Biorad) and a ChemiDoc ™ XRS Imaging system (BioRad).

## ELISA

The presence of OSM cytokine in the media of BV-2 cells was detected by solid-phase sandwich ELISA (R&D, Quantikine mouse OSM kit) with recommendations of the manufacturer. Absorbance measuring was done at 450 nm with a CLARIOSTAR microplate reader. 50 µl of media conditioned by BV-2 cells for 3 days was used for these experiments. Pure OSM supplied with the kit was used for reference curve.

### Statistical analysis and countings

All experiments and stainings were performed at least three times and most of them were done four times. For cell countings, independent fields (number specified in the legends, at least 6) were counted. Data are represented as means ± standard error of mean (S.E.M). Statistical differences in experiments were analyzed with tests indicated in the legends using GraphPad Prism software and BootstRatio website (http://rht.iconcologia.net/stats/br/) for QPCR (Clèries et al. 2012). Significances: *** (p<0.001), ** (p<0.01), * (p≤0.05).

### Equipment and settings

Fluorescent images were taken with a Zeiss apotome Axio Imager 2 equipped with a Zeiss ZEN software. Main settings were: binning 2×2, apotome mode: 5. Quantifications were obtained either manually or through image captures and ImageJ software cell counting tool.

### Accession number

The raw data that support the findings of this study are openly available at the functional genomics data Gene Expression Omnibus (GEO: GSE149669).

## Results

### Ependymal cell RNA profile is highly modified by SCI

We explored the mechanisms underlying ependymal region activation by performing moderate spinal cord injury in six mice. Six other mice with sham operation were used as control. After 3 days, the central canal region was microdissected (Fig. 1A, B) and RNAs were analysed with microarrays as previously described (Ghazale et al. 2019). Two distinct RNA profiles were observed in ependymal cells microdissected from control and injured spinal cord (Fig. S1A). Data analysis revealed that 510 genes were significantly upregulated (p ≤ 0.05; fold change ≥ 2) while 458 genes were downregulated (fold change ≤ −2) (Fig. 1C). Top upregulated and downregulated genes are shown in Table 1 and the full gene list is in Table S2. Six genes were upregulated more than 20 fold, namely *Crym, Ecm1, Ifi202b, Nupr1, Osmr, Rbp1, Thbs2* whereas only one was downregulated to this extent (*Acta1*). CRYM and RBP1 are two proteins implicated in pipecolic acid and retinol metabolism respectively (Siegenthaler 1996; Hallen et al. 2015). NUPR1 and IFI202B are involved in transcriptional response to various cellular stress notably by binding to P53 and by inhibiting apoptosis (Xin et al. 2006; Siegenthaler 1996). THBS2 is an adhesive glycoprotein and ECM1 is a protein of the extracellular matrix involved in angiogenesis (Sercu, Zhang, et Merregaert 2008). OSMR is a receptor for various IL-6 related cytokines (Morikawa 2005) and ACTA1 is an isoform of actin found in muscle tissues and which is highly enriched in ependymal cells (Ghazale et al. 2019). These modifications suggest that SCI trigger major changes in ependymal cell metabolism, transcriptional networks and phenotype.

We then submitted the dysregulated gene list to GO and pathways analyses (Fig. 1D and Table S2). As expected from previous works showing proliferation of ependymal cells after SCI, many genes related to cell cycle such as *Myc* and *Cyclins* (A2, B1/2 and D1/2) were upregulated by SCI. More interestingly, RNAs for several extracellular matrix genes such as Fibronectin (*Fn*), Matrilin (*Matn2*), Vitronectin (*Vtn*), Versican (*Vcan*), and several Collagens (*Col8A1, Col11A1, Col12A1, Col14A1*) were increased. Finally, ependymal cell reacted to SCI by the modification of expression of 63 genes closely related to transcription (42 and 21 up and down regulated respectively) (Table 1 and Table S2). Five were upregulated more than 10 fold (*Cebpd, Etv5, Fos, Nupr1, Ifi202b)*. CEBPD is an important transcription factor regulating the expression of genes involved in immune and inflammatory responses (Balamurugan et Sterneck 2013). ETV5 (also known as ERM) is a member of the PEA3 subfamily of ETS transcription factors. Its expression is notably controlled by the ERK/MAPK pathway and is often correlated to cell proliferation and migration (Li et al. 2012). Finally, FOS is a well-known proto-oncogene regulating cell proliferation and differentiation in many contexts (Kovács 1998).

To see whether these RNA variations led to protein level modifications, we selected two genes: one which was highly overexpressed after SCI (30x): *Osmr* the receptor for the oncostatin cytokine and another gene which was only moderately increased (2.5x), *Ntn* coding for NETRIN, a secreted protein involved in many processes such as stem cell renewal and apoptosis (Lai Wing Sun, Correia, et Kennedy 2011). Figure 2 shows weak or no detection of these 2 proteins by immunofluorescence (IF) in the control ependymal region but a clear induction in ependymal cells after injury. This shows that the SCI-induced transcriptional modifications lead to new proteins in ependymal cells.

To understand further the signaling pathways affected by SCI in the ependyma region, we used a causal analysis approach with Ingenuity pathway application (Krämer et al. 2014). Two pathways were identified (Fig. S1B): MAPK/ERK and STAT3 signalings. We validated these activations by performing IF against the phosphorylated forms of STAT3 and ERK/MAPK1 (p-STAT3 and p-ERK). Figure 2 shows that whereas we could not detect positive cells in control non-injured animals, a fraction of cells around the central canal expressed p-STAT3 and p-ERK after SCI. We decided to support our results further by using organotypic cultures of adult spinal cord slices (Fig. 1A). In this ex vivo model, culture conditions spontaneously activate ependymal cells which then proliferate (Fernandez-Zafra, Codeluppi, et Uhlén 2017). We performed IF for p-ERK, p-STAT3 on slices after 0 and 3 days of culture. Figure 3 shows that in this model, the p-ERK protein was clearly induced in a fraction of ependymal cells after 3 days of culture. No convincing and reproducible stainings were observed for p-STAT3.

**Figure 3:**
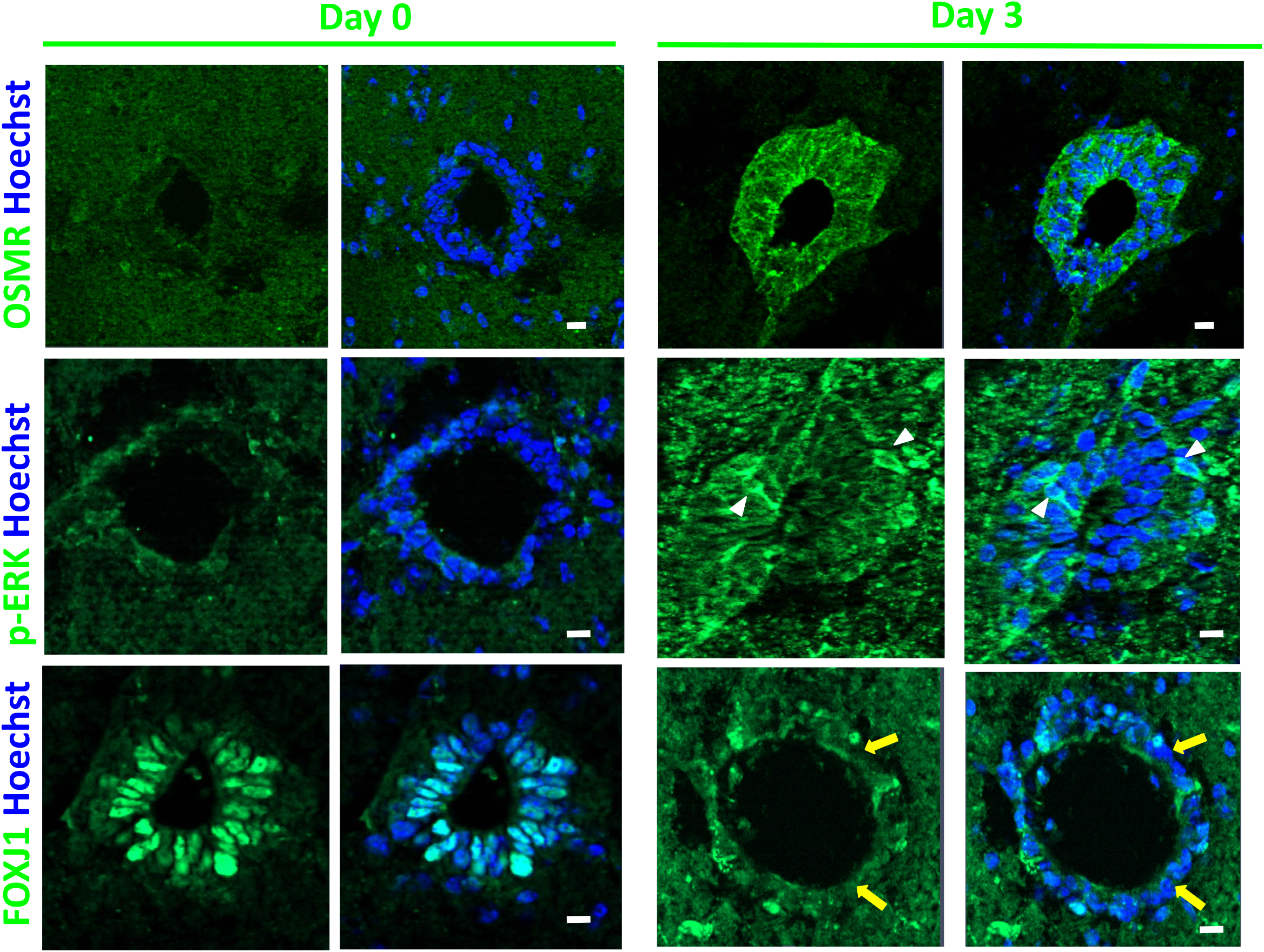
Spinal cord organotypic slice cultures: Immunofluorescences for indicated proteins performed on spinal cord slices after collection (Day 0) and after 3 days in culture (Day 3). White arrowheads and yellow arrow show examples of positive and negative cells respectively. These images are representative of 3 independent experiments. (n=4 animals, 5 sections examined per culture). Scale bars=10 µm.

### Cilia genes are downregulated after SCI and are regulated by growth factors

We previously reported that, consistent with their functions, ependymal cells highly expressed genes involved in ciliogenesis (Ghazale et al. 2019). Remarkably, GO analysis revealed the significant downregulation of these genes after SCI (Fig. 1D and Table S2). To illustrate this, we selected 12 genes (*Armc4, Ccdc151, Ccdc39, Ccdc40, Crocc, Dnah11, Lrrc6, Mak, Pacrg, Spef2, Stk36, Wdr19)* with documented role in cilia and which are very specifically expressed in the spinal cord ependymal region (fold change>16 compared to the spinal cord parenchyma, Ghazale et al. 2019). Microarray analyses show that the expression of these 12 genes were significantly downregulated between 1.5 and 2.7 fold after injury (Fig. 4A). This suggested that transcription factors governing ciliogenesis might be also affected by SCI. As we previously found (Ghazale et al. 2019) that mouse spinal cord ependymal cells specifically express several transcription factors involved in cilia formation, namely *FoxJ1, Rfx1-4* (Lemeille et al. 2020), *and Trp73* (Nemajerova et Moll 2019), we analysed their expression after SCI. Microarray results presented in Fig. 4A indicated that *Rfx1 and Trp73* RNA levels were significantly reduced whereas *Rfx4* was increased. With regards to *FoxJ1* and *Rfx2*, their expression also appeared to be reduced after SCI but did not reach significance (*FoxJ1:* mean +/-S.E.M.= 1.01+/-0.14 vs 0.86+/-0.22 post SCI and *Rfx2*: 1.01+/-0.21 and 0.76 +/-0.32 post SCI, n=6). As FOXJ1 is the central transcription factor orchestrating cilium morphogenesis (Lewis et Stracker 2020), we also explored its expression at the protein level by IF. Figure 2 shows that compared to control spinal cord, a reduction of FOXJ1 is observed in ependymal cells after SCI and some of them became negative. We validated this downregulation further using the spinal cord organotypic slice model mentioned previously (Fig. 1A). Figure 3 shows that, compared to control slices, FOXJ1 expression was barely detected in ependymal cells maintained for 3 days in culture.

**Figure 4:**
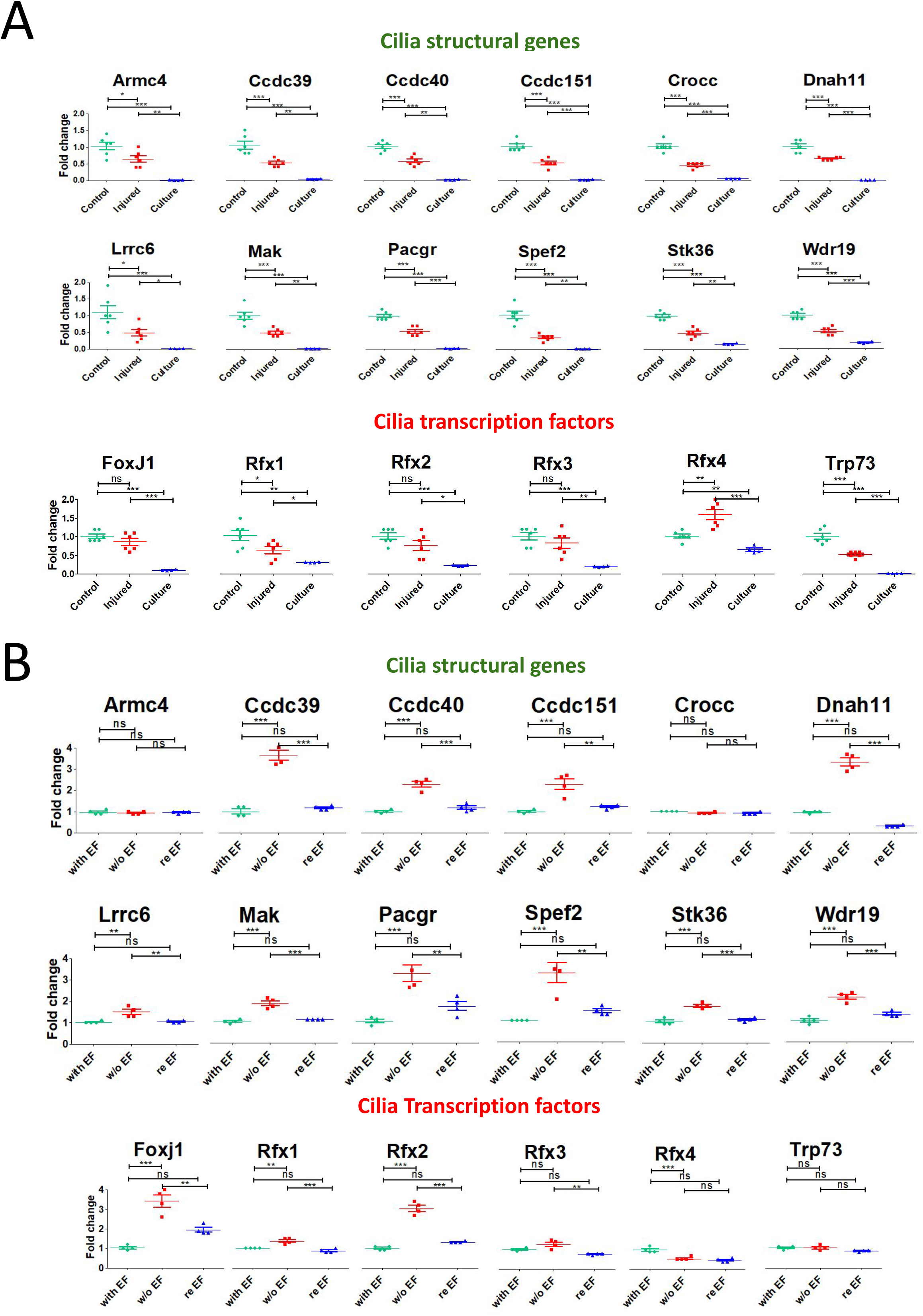
Cilia gene expression is affected by SCI and culture conditions. A:Microarray quantifications of indicated genes in 1/control spinal cord, 2/injured spinal cord, 3/ growing spinal cord neurospheres (n=6 for spinal cords and n=4 for neurosphere cultures). Tests=one way ANOVA with Tukey post tests. Values represent fold changes compared to control spinal cords. **B:** Microarray quantifications of indicated genes in neurospheres cultured in 3 conditions (n=4 for each conditions): 1/ with EF (i.e. growing with EGF and FGF2), 2/ w/o EF (i.e. differentiated by removing EGF and FGF2), 3/ re EF (reintroduction of EGF and FGF2 for 3 days after the differentiation step). Tests=one way ANOVA with Tukey post tests. Values represent fold change compared to neurospheres in the growing condition. n.s.=not significant

Next, we questioned the level of expression of these cilia-related genes in primary spinal cord neurospheres which mainly originate from the central canal ependymal cells (Meletis et al. 2008). We derived four independent neurosphere primary cultures from adult spinal cords and performed RNA profiling of these cells using microarrays. Data presented in figure 4A indicate that neurosphere cells expressed the 12 selected cilia genes and six cilia transcription factors (*FoxJ1, Rfx1-4, Trp73)* at a much lower level than in vivo ependymal cells.

Compared to ependymal cells which proliferate at a very low rate, neurophere cell are highly proliferative with a doubling time of approximately 28 hours (unpublished data). Considering the low expression of cilia genes found in neuropheres (Fig. 4A), we reasoned that these genes might be negatively controlled by growth factors. We tested this possibility by removing the growth factors (EGF/FGF2) in the media for 3 days and then by monitoring gene expression with microarrays (n=4). Indeed, Figure 4B shows that growth factor removal significantly increase the expression of 12/14 of cilia genes (fold change between 1.5 to 3.9) together with 3 transcription factors (*FoxJ1, Rfx1, Rfx2*) (fold change 3.4, 1.3, 3.1 respectively). Conversely *Rfx4* was downregulated. We validated this result at the protein level for FOXJ1. Indeed, whereas FOXJ1 is not detected by immunofluorescence in neurosphere cells cultured with growth factors, it is well-expressed after growth factor removal (Fig. 5A). To test further the influence of growth factors on the expression of these cilia genes, we reintroduced them in the media for 3 days and performed new microarray analysis (n=4). This led to a significant downregulation of 12/14 cilia genes together with *FoxJ1 Rfx1, Rfx2*. This illustrates the negative influence of growth factors on cilia genes and cilia transcription factors in neurosphere cells. Collectively, these results support the notion that SCI opposes cilia gene expression in ependymal cells, possibly by the activation of proliferation and the downregulation of FOXJ1 transcription factor.

**Figure 5:**
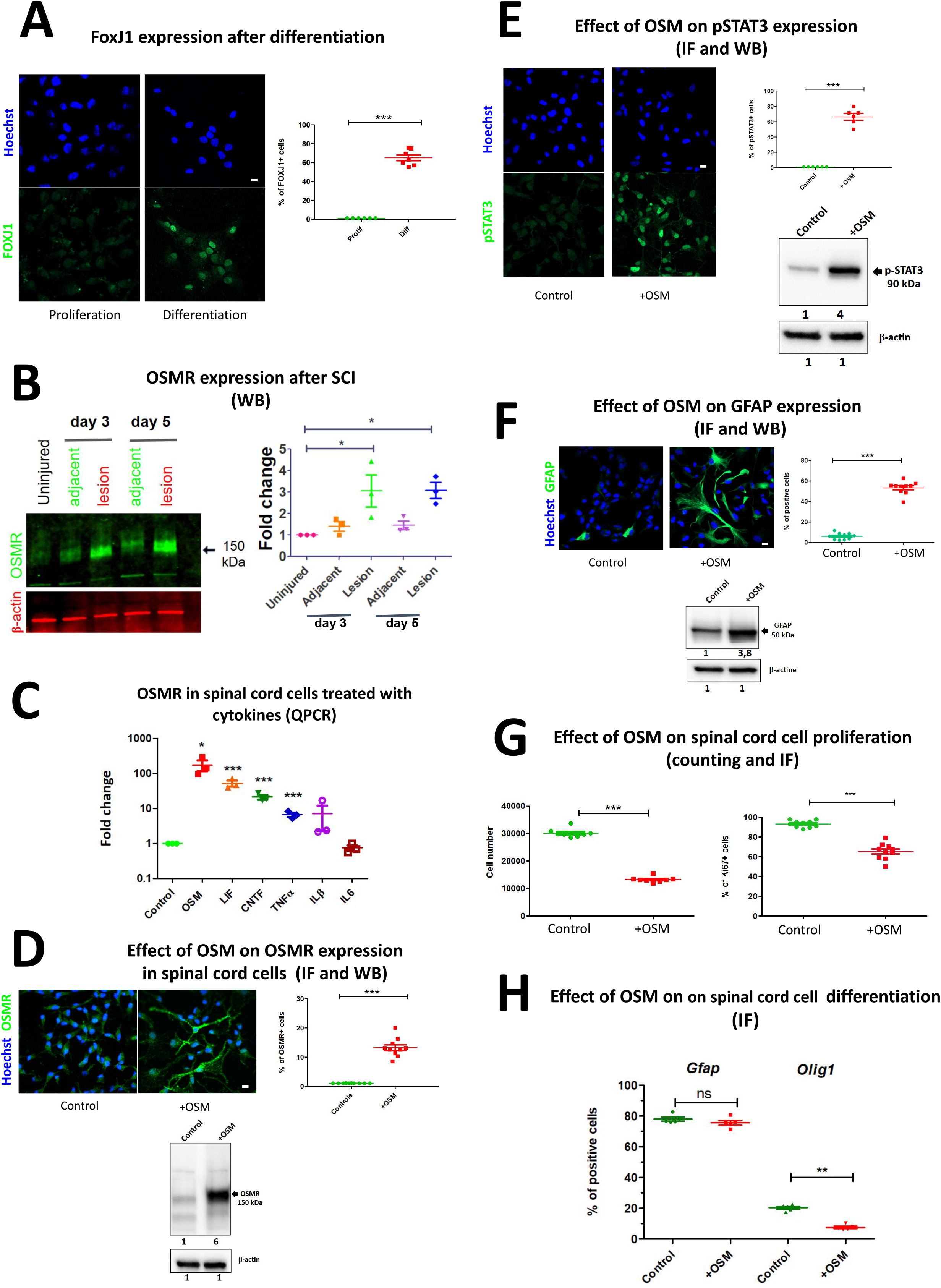
OSM affects OSMR expression, neurosphere growth and differentiation. A: *Left-hand images:* immunofluorescences for FOXJ1 in spinal cord stem cells cultured in proliferation (with growth factors) or differentiation (without growth factors) conditions. Scale bar=10 µm. *Right panels:* immunofluorescence quantification (n=7 fields). **B:** *Left-hand panel:* WB for OSMR on proteins extracted from control and injured spinal cord (3 and 5 days post injury). Proteins were extracted from spinal cord fragments containing and adjacent to the lesion site. β-actin was used for normalization. *Right-hand panel:* WB quantification. Values represent fold change compared to uninjured spinal cord protein extracts. Tests=one way ANOVA with Tukey post tests. n=3 independent experiments. **C:** QPCR for *Osmr* RNA in growing spinal cord neurospheres treated for 3 days with indicated cytokines. Values represent fold change compared to untreated neurospheres. Statistical tests were performed with Bootstratio (Clèries et al. 2012) compared to non-treated neurospheres. n=3 independent experiments. **D:** *Left-hand panels*: immunofluorescences for OSMR in control or OSM-treated spinal cord stem cells. Scale-bar=10 µm. *Right-hand panel*: immunofluorescence quantification (n=10 fields). Test= two tailed t-test. *Lower panel*. WB for OSMR on proteins extracted from untreated and OSM-treated neurospheres. Numbers represent quantification (fold change) compared to untreated neurospheres. β-actin was used for normalization. **E:** *Left-hand panels*: immunofluorescences for p-STAT3 (phospho-STAT3) in control or OSM-treated spinal cord stem cells. Scale-bar=10 µm. *Right-hand panel*: immunofluorescence quantification (n=6 fields). Test= two tailed t-test. *Lower panel*. WB for p-STAT3 on proteins extracted from untreated and OSM-treated neurospheres. Numbers represent quantification (fold change) compared to untreated neurospheres. β-actin is used for normalization. **F:** *Left panels:* Immunofluorescence for GFAP in untreated and OSM-treated cell cultures. *Right-hand panel*: immunofluorescence quantification (n=10 fields). Test= two tailed t-test. *Lower panel:* WB for GFAP on proteins extracted from untreated and OSM-treated neurospheres. Numbers represent quantification (fold change) compared to untreated neurospheres. β-actin is used for normalization. **G:** Effect of OSM on neurosphere growth. *Left-hand panel:* Diagram shows the number of cells obtained in untreated and OSM-treated neurospheres 6 days after seeding (n=8 wells). Test=two tailed t-test. *Right-hand panel:* Diagram shows the % of MKI67^+^ cells obtained in untreated and OSM-treated cell cultures (n=10 fields). Test=two tailed t-test. **H:** Effect of OSM on spinal cord stem cell differentiation. Diagram shows the % of GFAP^+^ and OLIG1^+^ cells, 4 days after differentiation in untreated and OSM-treated cell cultures. Tests=Mann-Whitney tests, n=5 independent experiments. n.s.=not significant

### Oncostatin affects proliferation and fate of spinal cord neurosphere cells

One gene we saliently found induced by SCI in ependymal cells is *Osmr*, a member of the IL-6 receptor family. Very little is known about this receptor in adult neural stem cells. This prompted us to investigate its role and regulation in the context of spinal cord ependyma. We started by validating the sharp upregulation of OSMR protein using the spinal cord organotypic slice model described above (Fig. 1A). Figure 3 shows that after 3 days of culture, ependymal cells highly expressed OSMR detected by IF. We also performed western blot with proteins extracted from uninjured spinal cord and 3 and 5 days after SCI. This again revealed a sharp induction of OSMR by SCI (Fig. 5B). It is worth noting that here, the detection of OSMR reflects its induction in ependymal cells but probably also in other undefined cells in the spinal cord parenchyma which may overexpress OSMR after SCI.

We then explored how OSMR is upregulated after SCI in ependymal cells. It has been shown that *Omsr* is induced by various cytokines, notably belonging to the IL-6 family, in different cell types. We evaluated this possibility in the context of the spinal cord using neurosphere cultures (Fig. 1A). These were treated for 5 days with Il-6 related cytokines (CNTF, Il-6, LIF, OSM), inflammatory cytokines (Interferon γ, CCL2, CXCL3, TGFb, TNFa) and various cytokines which have been shown to influence neural precursor fate (GDF15, VEGFC, BMP6). The effect on *Osmr* expression was measured by QPCR. We found that only four of these cytokines (CNTF, LIF, OSM and TNFa) upregulated *Osmr* expression (Fig. 5C). OSM, LIF and CNTF were particularly potent with a 175, 50 and and 20 increase respectively (Fig. 5C). Western blot and IF also validated the strong increase of OSMR after OSM treatment (Fig. 5D). In the brain, these Il-6 related cytokines can activate STAT3 signaling in neural stem cells which is accompanied by proliferation reduction and astrocytic differentiation (Taga et Fukuda 2005). Indeed we found that addition of OSM to growing neurospheres induced a sharp increase of p-STAT3 expression (Fig. 5E) and astrocytic differentiation as revealed by IF and western blot for GFAP (Fig. 5F). OSM also negatively affected neurosphere growth as measured by the reduced number of cells obtained 7 days after seeding and by the decrease in MKi67^+^ cells (Fig. 5G).

Lastly, we determined the influence of OSM on the differentiation of neurosphere cells into different cell types. Growing neurospheres were treated with OSM for 3 days and then for 3 additional days during the differentiation phase with serum. Differentiation into astrocytes, oligodendrocytes and neurons was quantified by immunostainings for GFAP, OLIG1 and MAP2/TUBB3. In control condition, most neurosphere cells differentiated into astrocytes (approximately 80%) and no significant effect of OSM was observed on their production (Fig. 5H). In contrast, OSM reduced by half the % of OLIG1+ cells indicating that this cytokine negatively impact on the formation of oligodendrocytic cells (Fig. 5H). With regard to the neuronal differentiation, the formation of MAP2^*+*^ or TUBB3^*+*^ cells was very weak (typically under 1%) and highly variable between experiments and thus no reliable conclusion could be obtained.

All together, these results indicate that spinal cord neurosphere cells are highly responsive to OSM cytokine which alters their proliferation and fate.

### OSM expression is strongly upregulated in SCI

Considering the major effect of OSM on OSMR expression, proliferation and fate of spinal cord cells in vitro, we questioned the origin of this cytokine. Two brain single cell RNA databases indicated that *Osm* is almost exclusively expressed by microglia cells especially when they are activated (Fig. S2A and B). To see how *Osm* expression varies during SCI, we performed QPCR on spinal cord RNA extracted before and 3 days post injury (n=3 mice). We found that *Osm* is strongly upregulated after SCI (Fig. 6A) (fold change: 5-10x, n=3 mice). Expression of *Cntf, Lif and Tnfa*, which we showed can upregulate *Osmr* in neurosphere cells (Fig. 5C), are also increased in spinal cord after injury but more moderately than *Osm* (Fig. 6A). These striking upregulations of *Osm, Lif* and *Cntf* are also observed in a database comparing gene expression before and after spinal cord injury (Fig. S2C) (Chen et al. 2013).

**Figure 6:**
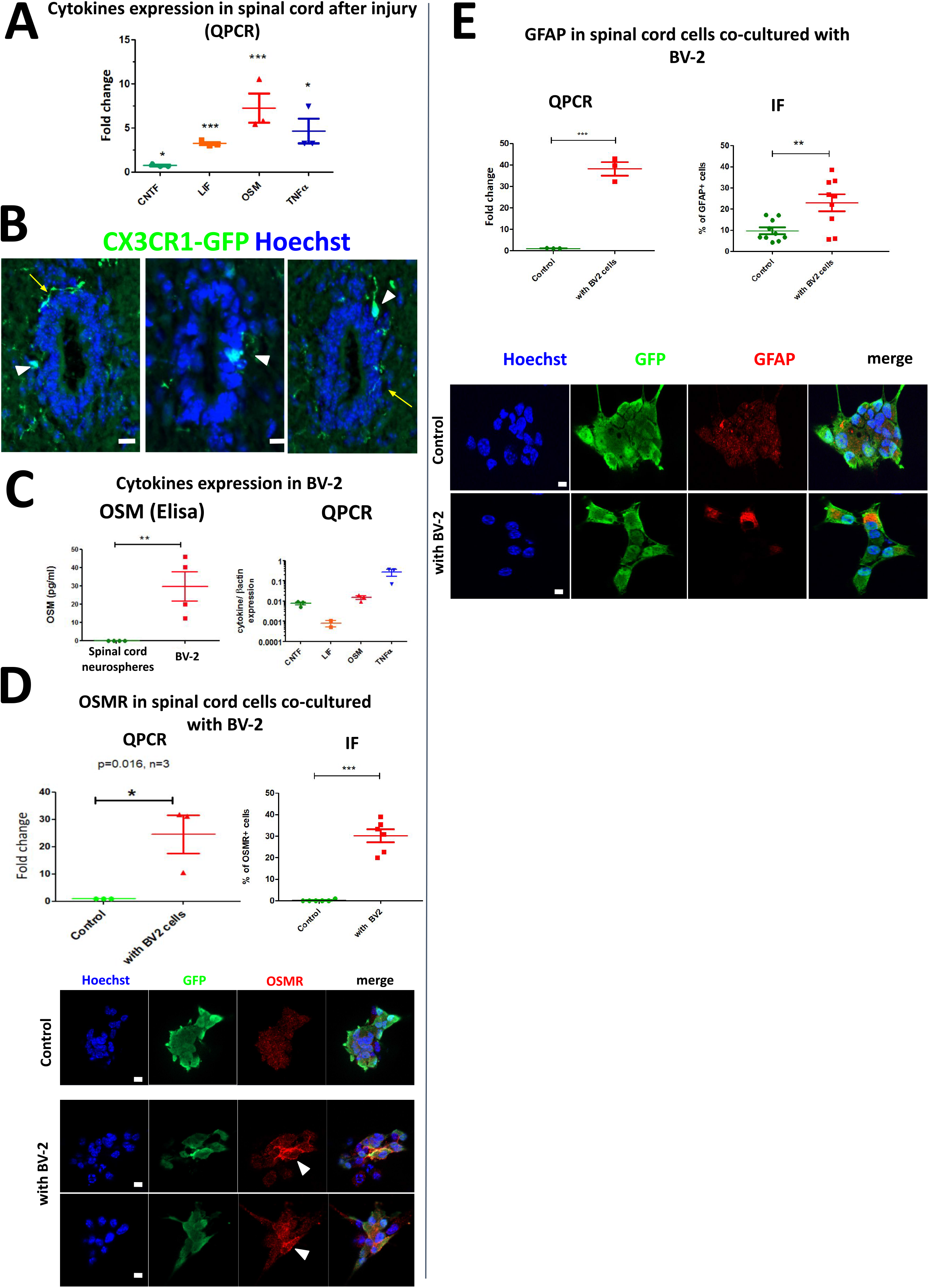
Microglia cells upregulate OSMR expression in spinal cord stem cells. A QPCR for indicated cytokines in RNA extracted in sham-operated and injured spinal cord (3 days after SCI). n=3 independent experiments. Numbers indicate the fold change compared to sham-operated spinal cords. Statistical tests were performed with Bootstratio (Clèries et al. 2012) compared to control spinal cords. **B:** Immunofluorescences for GFP performed in CX3CR1-GFP mice to reveal microglial cells (green) associated to ependymal cells. White arrowheads and yellow arrows show microglia cell somas and processes respectively, close or within the ependymal cell layer. Scale bars=10 µm. **C:** *Left-hand panel:* detection of the OSM cytokine in the supernatant of BV-2 microglial cell culture by ELISA (n=4 independent experiments). *Right-hand panel:* detection of indicated cytokines in BV-2 microglial cells by QPCR (n=3 independent experiments). **D:** Influence of BV-2 cells on OSMR expression in spinal cord stem cells. *Left-hand panel:* QPCR for OSMR. RNA were extracted from spinal cord neurosphere cells cultured without (control) and with BV-2 cells. Values represent fold change compared to control neurospheres. Statistical test was performed with Bootstratio (Clèries et al. 2012) compared to control neurospheres. n=3 independent experiments. *Lower panel:* Immunofluorescences for OSMR in GFP^+^ spinal cord stem cells cultured without (control) or with BV-2 microglial cells. Scale bars=10 µm. *Right-hand panel:* Quantification of immunofluorescences (n=6 fields). test= two tailed t-test. **E:** Influence of BV-2 cells on GFAP expression in spinal cord stem cells. *Left-hand panel:* QPCR for GFAP. RNA were extracted from spinal cord neurosphere cells cultured without (control) and with BV-2 cells. Values represent fold change compared to control neurospheres. Statistical test was performed with Bootstratio (Clèries et al. 2012) compared to control neurospheres. n=3 independent experiments. *Lower panel:* Immunofluorescences for GFAP in GFP^+^ spinal cord stem cells cultured without (control) or with BV-2 microglial cells. Scale bars=10 µm. *Right-hand panel:* Quantification of immunofluorescences (n=10 fields). test= two tailed t-test.

### Microglial cells upregulate Osmr in spinal cord neurosphere cells

Close interactions of microglial cells with neural stem cell have been described in the brain (Matarredona, Talaverón, et Pastor 2018). This prompted us to examine this possibility in the context of the spinal cord ependymal region. We used the CX3CR1-GFP transgenic mice to visualize microglial cells including their soma but also their processes as these cells are highly ramified. Figure 6B shows the presence of microglial cell soma (white arrowheads) in close contact with ependymal cells. Closer examinations also revealed microglial cell processes in the proximity of ependymal cells suggesting possible interactions (Fig. 6B yellow arrows). Interactions between these two cell types can also be observed in the Gensat database (Heintz 2004) with two other microglial specific transgenic GFP mice (*Limd2* and *Csf2rb2)* (Fig. S3).

These observations encouraged us to investigate whether microglial cells could increase *Osmr* expression in spinal cord neurosphere cells. We based our approach on the mouse BV-2 cell line which is acknowledged as a reliable model for microglial cells (Blasi et al. 1990; Henn et al. 2009). By using ELISA test, we demonstrated that in comparison to spinal cord neurosphere cells, BV-2 cells secrete OSM (Fig. 6C left). QPCR analysis also revealed that BV-2 cells express genes for other cytokines (*Cntf, Lif* and *TNFa)* (Fig. 6C right). Next, we tested the influence of BV-2 cells on *Osmr* expression by co-culturing neurosphere cells with these cells for 3 days. For this experiment, we used spinal cord neurospheres derived from the spinal cord of β-actin GFP transgenic mice which enabled us to sort these cells after co-culture. *Osmr* level was measured by QPCR. Results presented on Fig. 6D showed a strong upregulation (>10 fold) of *Osmr* after co-culture with BV-2 cells. This was validated by IF, showing that OSMR is detected in a fraction of neurosphere cells after co-culture with BV-2 cells (Fig. 6D right). Using QPCR and IF, we also found that co-culturing with BV-2 cells strongly upregulate GFAP in neurosphere cells (Fig. 6E). Collectively these results support the notion that microglial cells participate in *Osmr* upregulation and astrocytic differentiation of spinal cord ependymal cells possibly through release of OSM.

## Discussion

In this work, we built up a new corpus of knowledge on spinal cord ependymal cells which are a source for new cells after lesion. The reaction of ependymal cells to SCI has been observed since 1962 (Adrian et Walker 1962), however very little was known on the underlying molecular mechanisms. We addressed this key issue by studying the RNA profiles of ependymal cells during SCI. Three main conclusions summarized in figure 7 can be drawn from this work.

**Figure 7:**
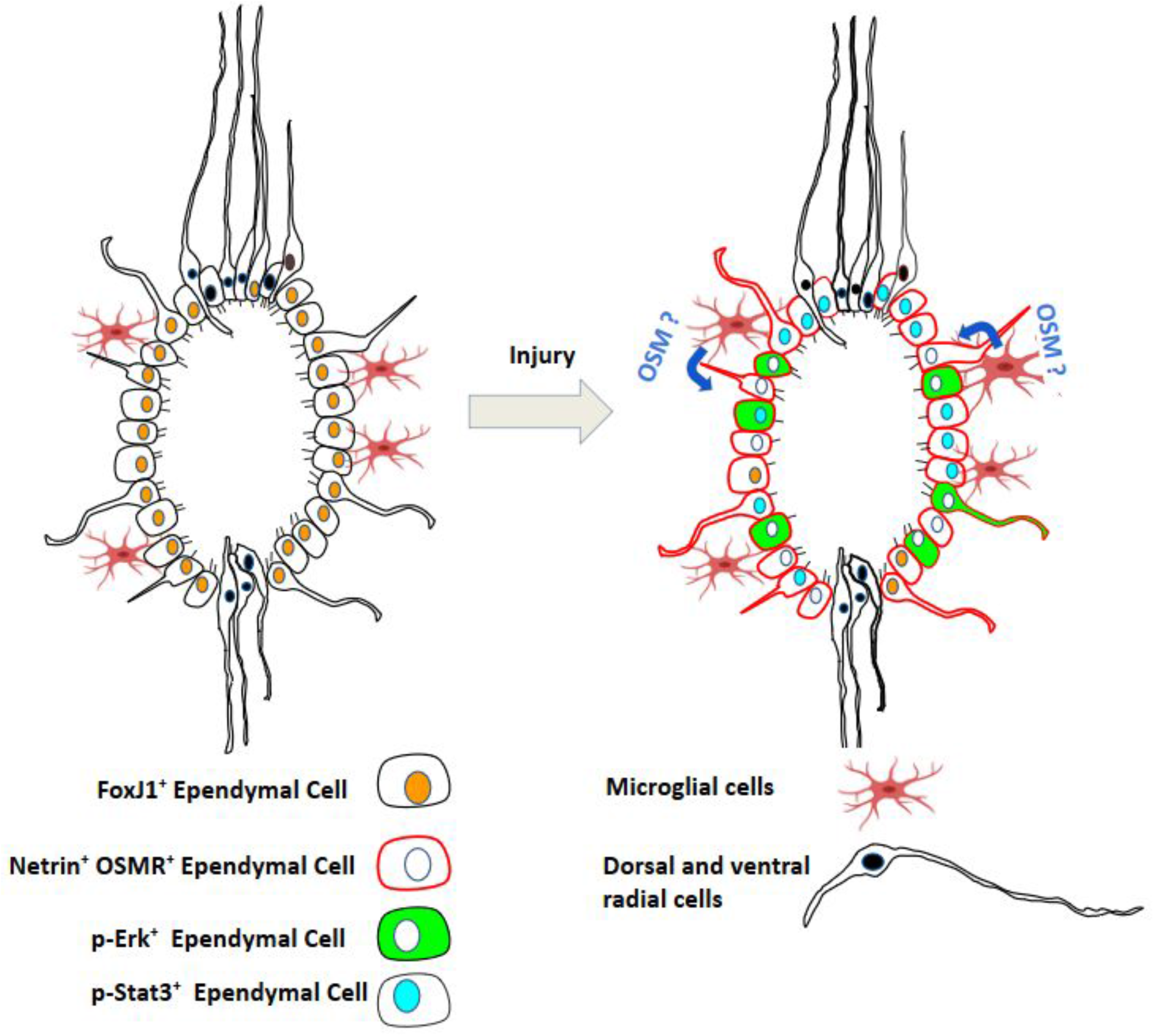
Graphical summary of main results. This schematic represents the different cell types found in the adult mouse spinal cord stem cell niche and their phenotypic modifications observed after injury. Dorsal and ventral radial cells have a distinct embryonic origin and express specific transcription factors compared to other ependymal cells (Becker, Becker, et Hugnot 2018). The behavior of these cells in SCI was not examined in this study so their representation is left unchanged in the schematic.

First, pathway analysis and IF demonstrated that SCI activates the ERK/MAPK and STAT3 pathways in a fraction of ependymal cells. What is the origin of these activations ? Many molecules such as IL-6 related cytokines (involved in inflammation), growth factors (i.e. EGF) and even ECM proteins such as fibronectin (Balanis et al. 2013) activate STAT3 and ERK/MAPK signalings. During SCI, the level of many of these proteins increases in the parenchyma (Chen et al. 2013) and in the ependyma itself (for instance fibronectin and the STAT-activating cytokine CLCF1 were upregulated 7x, Table S2). One possible candidate we identified is OSM but considering the variety of factors released during SCI, other molecules could contribute to activating STAT3 and ERK/MAPK pathways. What are the consequences of these activations on ependymal cells? Both pathways can promote astrocytic differentiation of brain neural stem cells (Li et al. 2012; Taga et Fukuda 2005). They may thus have a similar effect in spinal cord especially as GFAP is increased 6x in ependymal cells after SCI (Table S2). Besides differentiation and depending on their level of activation, these pathways can also promote stem cell proliferation (Niwa et al. 1998; Rhim et al. 2016). Thus they could also participate in the proliferation of ependymal cells induced by injury (Adrian et Walker 1962). ERK and STAT3 pathways could also upregulate several ECM genes in ependymal cells. For instance, expression of *Ecm1* and *Vcan* strongly increase after SCI (Table 1), and these genes are controlled by ERK and STAT3 in other contexts (He et al. 2018; Domenzain-Reyna et al. 2009). These ECM proteins may help migration of central canal cells toward the lesion site and/or fill extracellular space left by cellular death. Finally, activation of ERK and STAT3 pathways on ependymal cells likely act through downstream transcription factors. *Etv5* and *Cebpd* are potential candidates *as* their expression sharply rises after SCI (Table 1) and ERK and STAT3 pathways control their expression in other contexts (Li et al. 2012; Zhang et al. 2013).

A second significant result is that ependymal cells downregulate genes involved in ciliogenesis after SCI. In addition, whereas *Foxj1*, the master transcription factor of ciliogenesis, was not significantly decreased transcriptionally, we observed a reduction of the FOXJ1 protein in ependymal cells in vivo and also in vitro using spinal cord organotypic slices. This reduction could result from the degradation of FOXJ1 as this transcription factor has a short half-life and is constantly targeted by the ubiquitin-proteasome in brain ependymal cells (Abdi et al. 2018). Using primary spinal cord neurosphere cultures, we also observed that cilia genes and cilia transcription factors were expressed at a much lower level than in the spinal cord ependyma. Then, by removing and reintroducing growth factors in the media, we found that the expression level of a large fraction of these genes, notably *Foxj1*, increases and decreases respectively. These data are reminiscent of those obtained in brain ependymal cells (Abdi et al. 2018). Here, upon growth factor stimulation, brain ependymal cells lose *FoxJ1* expression and cilia, de-differentiate and proliferate. A similar scenario may apply in spinal cord ependymal cells in SCI. The mechanisms which suppress ciliogenesis in dividing cells have been partly elucidated in cancer and ERK signaling is involved (Higgins, Obaidi, et McMorrow 2019). Further investigations are needed to test whether ERK also impacts ciliogenesis in ependymal cells during injury.

Finally, this work unveiled the OSM/OSMR pathway as part of the reaction of spinal cord ependymal cell to SCI. OSMR was the second top upregulated gene after SCI with a 30 fold increase, which we confirmed at the protein level. Using in vitro experiments, we identified this pathway as a regulator of spinal cord cell proliferation and fate. Indeed OSM reduced neurosphere cell proliferation and increased p-STAT3, OSMR and GFAP. OSM also reduced the fraction of OLIG1^+^ oligodendrocytic cells during differentiation. These results are consistent with the role of OSM/OSMR in other contexts. In brain, OSM reduces the proliferation of neural stem cell in vitro and their number in vivo (Beatus et al. 2011) while pushing toward astrocytic differentiation (Yanagisawa, Nakashima, et Taga 1999). Besides, in various models OSM can induce both p-STAT3 and SOCS3 (Stross et al. 2006), a protein limiting STAT3 signaling and which is one of the top 10 gene upregulated after SCI (Table 1).

What triggers *Osmr* expression in ependymal cells after SCI? We found that OSM is a very potent inducer of *Osmr* expression in neurosphere cells. In vivo, *Osm* gene expression is rapidly and strongly increased after injury in brain (x20, (Oliva et al. 2012)) and in spinal cord (x100 (Slaets et al. 2014), x20 (Chen et al. 2013), x5-10 this study). Transcriptomics databases revealed that *Osm* is almost exclusively expressed by microglial cells and we found that OSM-producing microglia cells can upregulate OSMR in spinal cord neurospheres in vitro. In addition, we observed close interactions of microglial cells with ependymal cells in vivo. Altogether, OSM derived from microglia cells appears as a strong candidate to orchestrate OSMR expression in ependymal cells. Microglia cells could contribute to the astrocyte differentiation of spinal ependymal cells as found in the brain (Nakanishi et al. 2007; Zhu et al. 2008). Nevertheless, we also observed that OSMR can be triggered by other cytokines such as LIF or CNTF which are expressed by reactive astrocytes in SCI (Banner, Moayeri, et Patterson 1997) and literature shows that lipids such as prostaglandin E2 also increase OSMR (Ganesh et al. 2012). Thus, in addition to OSM and microglial cells, several molecules and reactive astrocytes may contribute to increasing OSMR in ependymal cells after injury which merits further investigation.

What is the significance of OSMR expression in spinal cord ependymal cells in vivo ? In vitro, we found that OSM influences spinal cord neurosphere differentiation toward astrocytes vs oligodendrocyte cells concomitantly with an increase in GFAP, OSMR and p-STAT3 expression. In the context of astrocytic differentiation of brain fetal progenitors, *Gfap* and *Osmr* genes are also coregulated by STAT3 signaling (Yanagisawa, Nakashima, et Taga 1999). Thus, the observed upregulation of OSMR and p-STAT3 in ependymal cells in vivo suggest their commitment toward astrocytic differentiation may be at the expense of oligodendrocyte lineage cells.

In conclusion, it remains unanswered why compared to zebrafish, mammal ependymal cells generate mainly astrocytes, few oligodendrocytes and no neurons after SCI. This could be due to astrocytic-fate determining factors expressed by these cells prior to injury such as SOX9 or NFIA transcription factors (Ghazale et al. 2019). In addition and reminiscent of the zebrafish situation (Kyritsis, Kizil, et Brand 2014), here we found evidence that their fate could be influenced by inflammatory cytokines provided by the environment, possibly microglia cells. Collectively, this work constitutes a molecular resource to study further the reaction of ependymal cell to SCI. This resource also sheds light on genes which are poorly annotated such as *Tchh, Rgs20, Gbp3* and which are highly-deregulated during SCI in ependymal cells. One limitation of this work is that considering the cellular heterogeneity present in the ependymal region (Ghazale et al. 2019), some identified gene variations may only apply to some cell subpopulations. This issue will be addressed thanks to the recent advances in single cell RNA seq analysis.

## Supporting information

Table 1 top genes

Table S1 primers and antibodies

Table S2 Gene analysis

## Acknowledgements

This work was supported by grants from IRP (Switzerland), IRME (France), AFM (France), ANR ERANET Neuroniche (JP Hugnot). H Ghazale was supported by an AFM PhD fellowship. We thank all Montpellier biocampus facilities for help (RHEM, MRI RIO, RAM, CECEMA) and excellent technical work. We are very grateful to Vicky Diakou (imaging), Dr C Duperray (cytometry), M Maistre (laser microdissection facilities, Bordeaux), V Pantesco (Affymetrix facilities) for providing technical expertise in this work.

The authors declare that there is no conflict of interest regarding the publication of this article.

## Author contributions

CR, HG, TC, FP, NH, DM, NG, CC and JPH performed and analyzed most of the experiments.

FP, HN, CC NG contributed to spinal cord injury models

CR, HG, TC, JPH contributed to histology, QPCR, WB, animal maintenance and cell culture

JPH wrote most the article which was edited by CR and HG

HH, AH provided reagents and mouse models for microglial cells as well as intellectual inputs.

