## supplemental figures for "Reaction of ependymal cells to spinal cord injury: a potential role for oncostatin pathway and microglial cells"

#### Supplemental figure legends

**Figure S1: Data analysis. A:** Heatmap of hierarchical clustering of genes differentially expressed in the ependymal region (control vs injured, p value ≤0.05, fold change≥2 or ≤-2, 1226 genes, n=6 mice per group). Sample numbers are indicated under the diagrams. **B:** Ingenuity pathway analysis of differentially expressed genes.

**Figure S2: OSM expression in normal and injured CNS. A:** In human and mouse CNS atlas (Brain RNA seq database Ben Barres's lab) OSM is specifically expressed in microglial cells. These images are derived from the database (brainrnaseq.org) associated to this publication (Zhang, et al. Purification and Characterization of Progenitor and Mature Human Astrocytes Reveals Transcriptional and Functional Differences with Mouse. Neuron (2016). **B**: *Upper panel*: Single cell RNA expression confirms that OSM is mainly expressed by immune cells (microglia and macrophage) in the mouse CNS. These data were obtained from the database (mousebrain.org/) associated to this publication (Zeisel et al, Cell. 2018 Aug 9;174(4):999-1014). Adult cell subtypes showing OSM expression are indicated by a blue rectangle under the hierarchical tree. Red arrow points to the microglia and macrophage subpopulations. *Lower panel*: Quantitative expression of OSM in these cells is presented as a blue circle and underneath value. **C:** Expression of *Cntf*, *Lif* and *Osm* cytokines by RNA seq after acute spinal cord injury. Data represent fold change compared to non injured spinal cord (day 0). These data were plotted after extraction from the database associated to this publication (RNA-seq characterization of spinal cord injury transcriptome in acute/subacute phases: a resource for understanding the pathology at the systems level, Chen K et al, PLoS One. 2013 Aug 9;8(8):e72567. doi: 10.1371/journal.pone.0072567)

**Figure S3:** Images imported from the Gensat expression database (gensat.org, Heintz N, Nat Neurosci. 2004 May;7(5):483) for 2 microglia specific genes (Limd2 and Csf2rb2). Brown stainings are IHC for GFP protein expressed under the control of Limd2 and Csf2rb2 promoters. Right-hand images are high magnification of the central canal region (cc) surrounding by the ependyma. Red arrows indicate microglial processes surrounding the ependymal region.

**FIGURE S1**

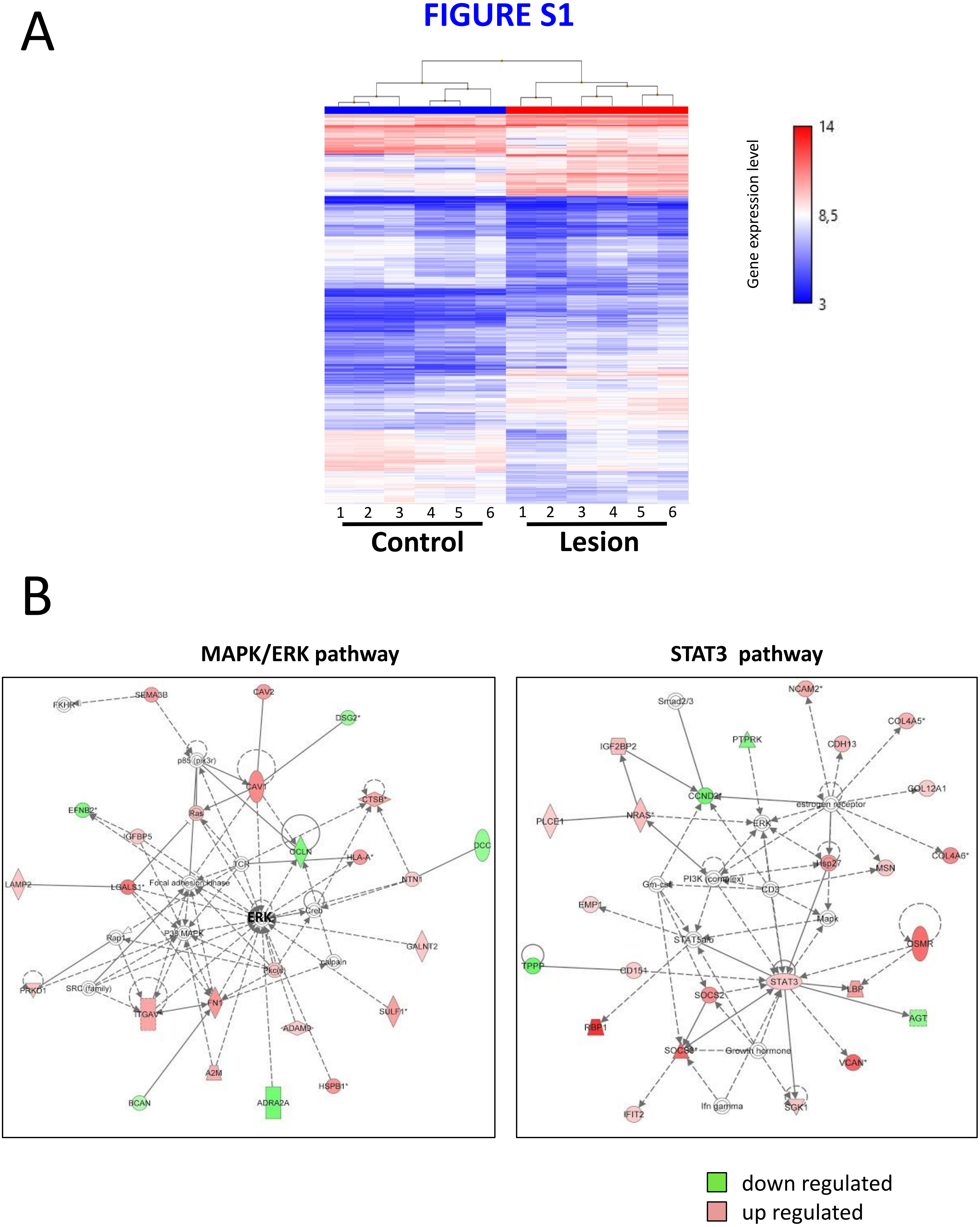
A
14
8,5
3
Gene expression level
1 2 3 4 5 6 1 2 3 4 5 6
Control
Lesion
B
MAPK/ERK pathway
STAT3 pathway
FKHR
SEMA3B
CAV2
DSG2*
p85 (pik3r)
CAV1
CTSB*
EFNB2*
Ras
IGFBP5
OCLN
DCC
HLA-A*
TCR
LAMP2
LGALS1*
Focal adhesion kinase
NTN1
ERK
Creb
Rap1
P38 MAPK
GALNT2
Pkc(s)
calpain
PRKD1
SRC (family)
ITGAV
FN1
SULF1*
ADAM9
A2M
HSPB1*
BCAN
ADRA2A
NCAM2*
Smad2/3
COL4A5*
IGF2BP2
PTPRK
CDH13
CCND2*
COL12A1
estrogen receptor
PLCE1
NRAS*
ERK
COL4A6*
Hsp27
MSN
Gm-csf
PI3K (complex)
CD3
EMP1
Mapk
STAT5a/b
OSMR
TPPP
CD151
STAT3
SOCS2
LBP
RBP1
AGT
SOCS3*
Growth hormone
VCAN*
IFIT2
Ifn gamma
SGK1
down regulated
up regulated

#### FIGURE S2

A

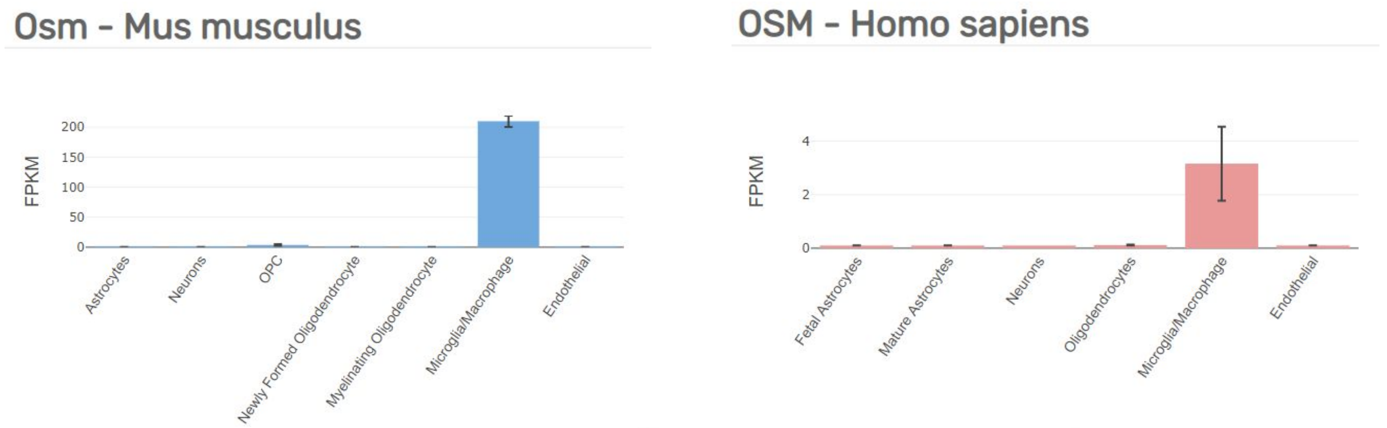


B

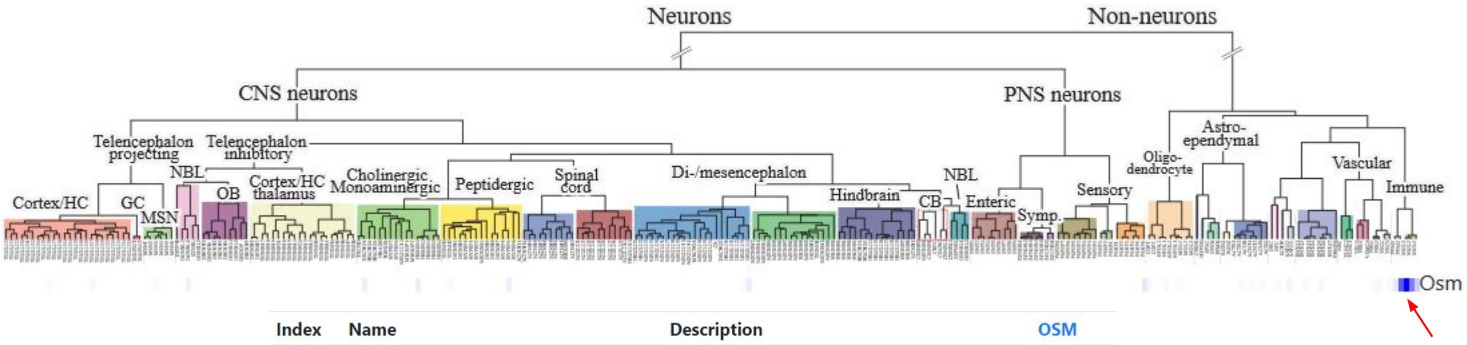


| Index | Name | Description | OSM |
|---|---|---|---|
| 261 | PVM1 | Perivascular macrophages | 0.0139 |
| 262 | PVM2 | Perivascular macrophages, activated | 0.135 |
| 263 | MGL3 | Microglia, activated | 0.210 |
| 264 | MGL2 | Microglia, activated | 0.104 |
| 265 | MGL1 | Microglia | 0.0381 |

C

**RNA levels after SCI**

(adapted from data published in Chen K et al, PLoS One. 2013 Aug 9;8(8):e72567)

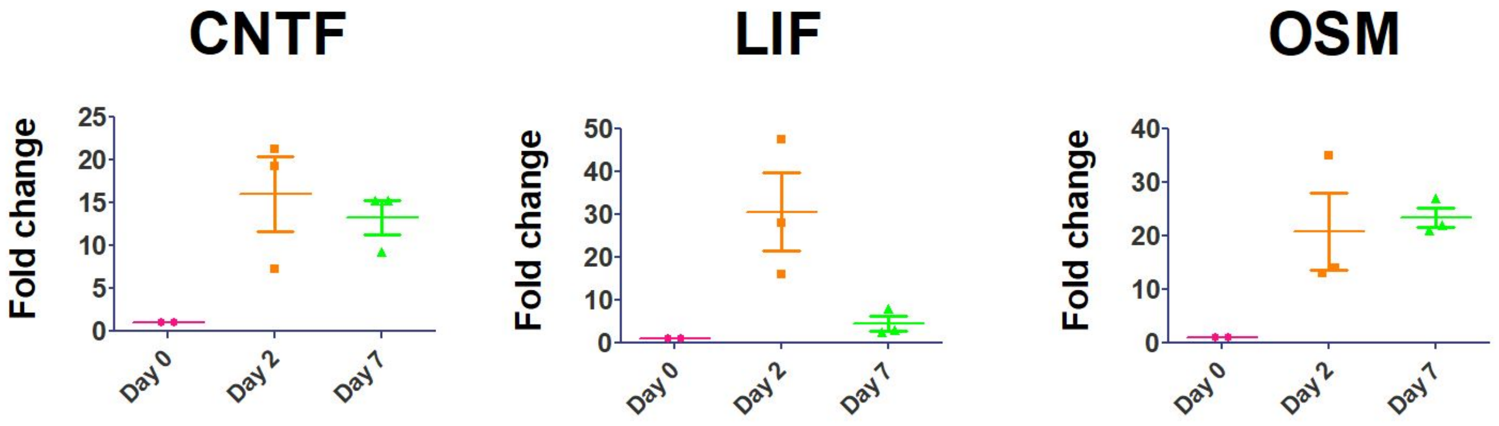

### FIGURE S3

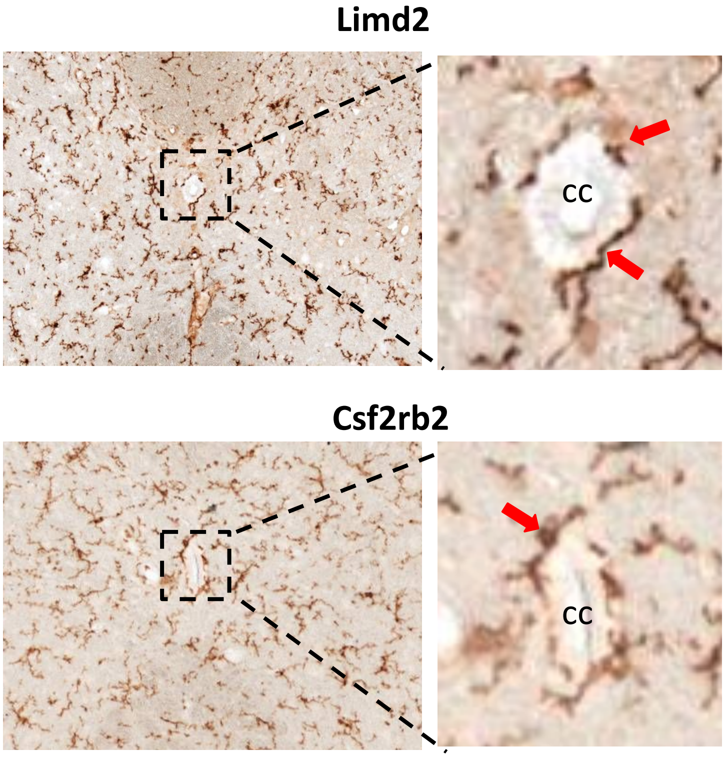
