## Supplementary material for "Reaction of ependymal cells to spinal cord injury: a potential role for oncostatin pathway and microglial cells": Table S1 primers and antibodies

**Antibodies**

| ***Name*** | ***Specie*** | ***Supplier*** | ***Reference*** | ***Dilution for IF*** | ***Dilution for WB*** |
| --- | --- | --- | --- | --- | --- |
| **β-ACTIN** | Mouse | Cell Signaling Technology | 8H10D10 | N.A. | 1/1000 |
| **FOXJ1** | Mouse | eBioscience | 14-9965-82 | 1/100 | N.A. |
| **GFAP** | Rabbit | Dako | Z0334 | 1/2000 | 1/5000 |
| **GFP** | Rabbit | Abcam | ab183734 | 1/500 | N.A. |
| **KI67** | Mouse | Pharmingen | 556003 | 1/500 |  |
| **NETRIN** | Rabbit | Abcam | ab126729 | 1/500 | N.A. |
| **OLIG1** | Goat | R&D | AF2417 | 1/100 | N.A. |
| **OSMR** | Goat | R&D | AF662-SP | 1/500 | 1/1000 |
| **p-ERK** | Rabbit | Cell Signaling Technology | 4370 | 1/200 | N.A. |
| **p-STAT3** | Rabbit | Cell Signaling Technology | #9131 | 1/500 | 1/250 |
| **TAG (**DYKDDDDK ) | Mouse | Supernatant | HB-9259 | 1/500 | N.A. |

**Primers**

| ***Name*** | ***Specie*** | ***Forward*** | ***Reverse*** | ***Size*** |
| --- | --- | --- | --- | --- |
| ***b-Actin*** | Mouse | GGCTGTATTCCCCTCCATCG | CCAGTTGGTAACAATGCCATGT | 154 |
| ***Cntf*** | Mouse | GGTGACTTCCATCAGGCAAT | CCATCAGCCTCTTTTTCAGG | 116 |
| ***Lif*** | Mouse | AGAAGGTCCTGAACCCCACT | CCACACGGTACTTGTTGCAC | 117 |
| ***Osm*** | Mouse | TCAGGGGTCTGATGACACAA | GAGAGAGCACCGGATCAGAC | 160 |
| ***Osmr*** | Mouse | CATCCCGAAGCGAAGTCTTGG | GGCTGGGACAGTCCATTCTAAA | 110 |
| ***Tnfa*** | Mouse | AGGGGATTATGGCTCAGGGT | GAGTCCTTGATGGTGGTGCA | 120 |
